# DNA Nanostructure-Guided Assembly of Proteins into Programmable Shapes

**DOI:** 10.1101/2023.08.07.552347

**Authors:** Qinyi Lu, Yang Xu, Erik Poppleton, Kun Zhou, Petr Sulc, Nicholas Stephanopoulos, Yonggang Ke

## Abstract

The development of methods to synthesize protein oligomers with precisely controlled number and configuration of components will enable artificial protein complexes and nanostructures with diverse biological and medical applications. Using DNA hybridization to link protein-DNA conjugates provides for a programmability that can be difficult to achieve with other methods, such as engineering multiple orthogonal protein-protein interfaces. However, it is still difficult to construct well-defined protein assemblies, and the challenge is magnified if only a single type of building block is used. To overcome this hurdle, we use a DNA origami as an “assembler” to guide the linking of protein-DNA conjugates using a series of oligonucleotide hybridization and displacement operations. We constructed several isomeric protein nanostructures on a DNA origami platform by using a C3-symmetric building block comprised of a protein trimer modified with DNA handles. By changing the number of protein-DNA building blocks attached to the origami, and the sequence of the linking and displacement strands added, we were able to produce dimers, two types of trimer structures, and three types of tetramer assemblies. Our approach expands the scope for the precise design and assembly of protein-based nanostructures, and will enable the formulation of functional protein complexes with stoichiometric and geometric control.

## Introduction

Cells are comprised of many modular supramolecular complexes, often made up of proteins, which perform diverse biological roles. The assembly of proteins into defined oligomeric structures allows for functions such as structural actuation in muscles^1^, transport along the membrane via ion channels^2^, and multienzyme catalysts^3^. Synthetic protein assemblies are therefore a promising class of biomaterials that can mimic, or potentially surpass, the uses of naturally occurring protein complexes^4, 5^. Proteins in such complexes are normally cohered by noncovalent protein–protein interactions^6, 7^, including hydrogen bonding, electrostatic interactions, van der Waals forces, or the hydrophobic effect^8, 9^. However, these protein-protein interactions require physical contacts between two or more molecules, with high specificity and a high degree of orthogonality. Although protein design approaches have showed great promise in engineering specific protein-protein interfaces (e.g., by using covalent “tags” form oligomeric complexes^10-14^, or *de novo* computational engineering of specific interfaces^15-21^, it is still difficult to design highly complex protein assemblies, especially from a common subset of building blocks.

One way to circumvent this limitation is to use protein-DNA conjugates, and link proteins into higher-order structures using programmable DNA linkers^22^. Since DNA strands with defined lengths and sequences are linked to the protein covalently, the specificity is mediated by the DNA strand (which affords hundreds or even thousands of sequence-defined orthogonal interactions^23, 24^) rather than the protein surface^25^, with the proteins subsequently linked to each other via DNA duplexes. Strategies for DNA-directed protein assembly can be generally divided to two categories. The first strategy places single-stranded (ss) DNA-modified proteins at designated positions on DNA nanostructures (using complementary DNA “docking” handles) to achieve controlled assembly^26-29^. In recent years, DNA origami^30-33^ has been widely used for organization of proteins with this approach, in which the origami nanostructure is an integral component of the final product. However, the massive size of DNA origami compared to individual proteins can limit the design and functions of the assembled protein complexes. The second strategy relies on connecting the oligonucleotide handles on proteins, either through attaching directly complementary strands to the proteins, or introducing additional DNA connectors. Although this approach can produce more compact protein assemblies with a limited amount of DNA, it requires the synthesis of site-specific protein-DNA conjugates. For relatively simple oligomers (e.g., dimers, linear trimers) or extended, periodic 1D^34, 35^ or 3D structures^36, 37, 38^, such synthesis is generally feasible. Nevertheless, the scalability of this strategy is questionable for design and assembly of larger and precise oligomers, since increasing the number of proteins and DNA strands will inevitably lead to incorrect assemblies. To avoid spurious nanostructure formation, and achieve more complex structures, proteins must be modified multiple, orthogonal DNA strands **(Figure 1A**), which in turn requires multiple site-specific reactions and purification of the desired conjugates. Such a task is not only synthetically challenging, but also decreases the yield of the final building blocks due to incomplete reaction and losses during purification.

**Figure 1:**
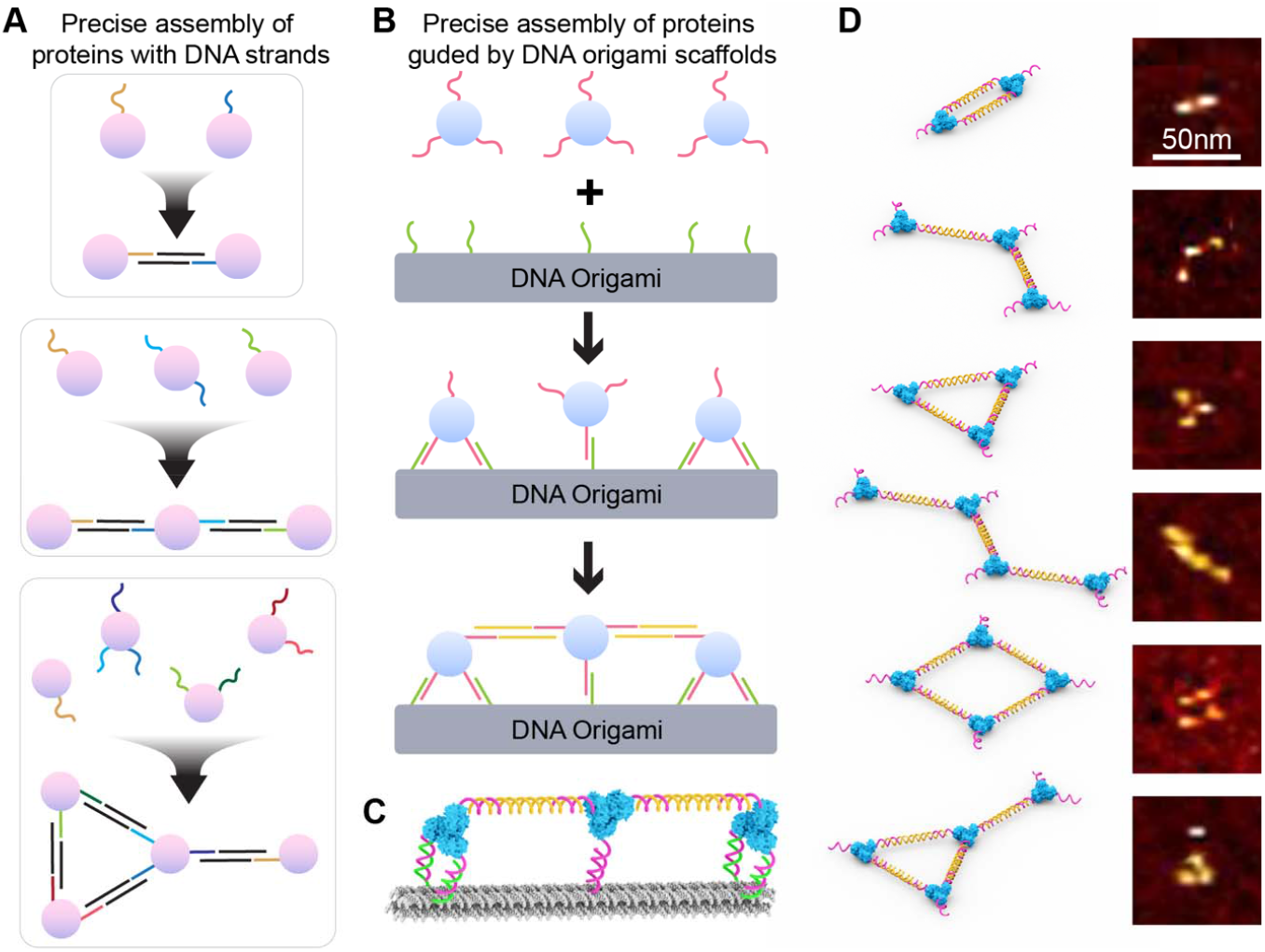
Protein-DNA conjugate connection strategies. (A) In the absence of a template, synthesizing a protein dimer, linear trimer, and Y-shaped tetramer requires proteins with multiple, orthogonal DNA handles, which is synthetically very challenging. (B) By contrast, a DNA origami scaffold—combined with precise addition of DNA linkers and displacement strands—can generate complex shapes from a single, oligomeric protein-DNA building block. (C) A 3D model of the origami-templated trimer in (B). (D) Protein oligomers produced in this work by using DNA origami templates (top-bottom): dimer, linear trimer, triangular trimer, linear tetramer, square-shaped tetramer, and “Y-shaped” tetramer.

To address aforementioned limitations in DNA-directed protein assembly, here we present a hybrid strategy that uses DNA origami as a nanoscale “assembler” to guide the assembly of proteins via a series of DNA anchoring, connecting, and displacement operations, forming a shape-defined protein structure (**Figure 1B**) that can be liberated from the origami. As a proof-of-concept, we used a homotrimeric protein-ssDNA conjugate based on a thermally stable C3 symmetric aldolase protein^39^ In the absence of a DNA origami template, the protein-ssDNA will form random-sized aggregates when mixed with a double-stranded DNA (dsDNA) connector. On the other hand, by using a combination of carefully designed anchor DNA strands, and controlling when and how the DNA handles on the protein are connected and displaced, precise protein oligomers can be assembled on, and then released from, a DNA origami structure (**Figure 1B and 1C**). Compared to previous methods, a distinctive feature of this strategy is that it can use the same protein-DNA building blocks and DNA origami template to construct different protein oligomers. The number, arrangement pattern, and distance between proteins can all be precisely designed by the number of docking sites on the origami, and the order in which the proteins are connected, which in turn makes it possible to control stepwise protein assembly into the desired shape. To demonstrate this versatility, we designed and tested different reaction routes for assembly of specific protein oligomers, including a dimer, linear trimer, triangular trimer, linear tetramer, square tetramer, and “Y-shaped” tetramer (**Figure 1D**).

## Results and Discussion

For the trimeric protein used as our building block, we selected 2-dehydro-3-deoxyphosphogluconate (KDPG) aldolase (ald), a 25 kDa protein that self-assembles into a C3-symmetric trimer (which we term ald_3_)^40, 41^. The symmetricity and size of ald_3_ (which can be approximated as a disk 6 nm diameter and 3 nm thickness) provide a promising building block for construction of nanostructures using DNA linkers. Although ald is an enzyme, here we used it exclusively as a structural building block due to its symmetry, high stability (> 80 °C), and ease of expression and modification with DNA. The homotrimeric nature of the ald_3_ oligomer also allows for multiple connections and more complex assemblies (compared with a monomeric or dimeric protein). Glutamate 54, a solvent-exposed residue on the outer edge of the trimer, was selected as an appropriate site for modification with DNA. We mutated this residue to cysteine (E54C) in order to perform thiol-selective chemistry with the heterobifunctional crosslinker succinimidyl 3-(2-pyridyldithio) propionate (SPDP) and a 5′-amino-modified oligonucleotide, as previously described^39^. After exposure to 6 equiv of purified succinimidyl 3-(2-pyridyldithio)propionate (SPDP) modified DNA, a band with higher retention was observed by electrophoresis on a denaturing polyacrylamide gel (SDS-PAGE), corresponding to oligonucleotide-bound ald monomer^39^. The yield is estimated to be ∼50% (**Supplementary Figure S1**). The ald_3_-DNA trimer (**Figure 2A**) was purified away from trimers bearing fewer DNA strands using anion exchange chromatography.

**Figure 2:**
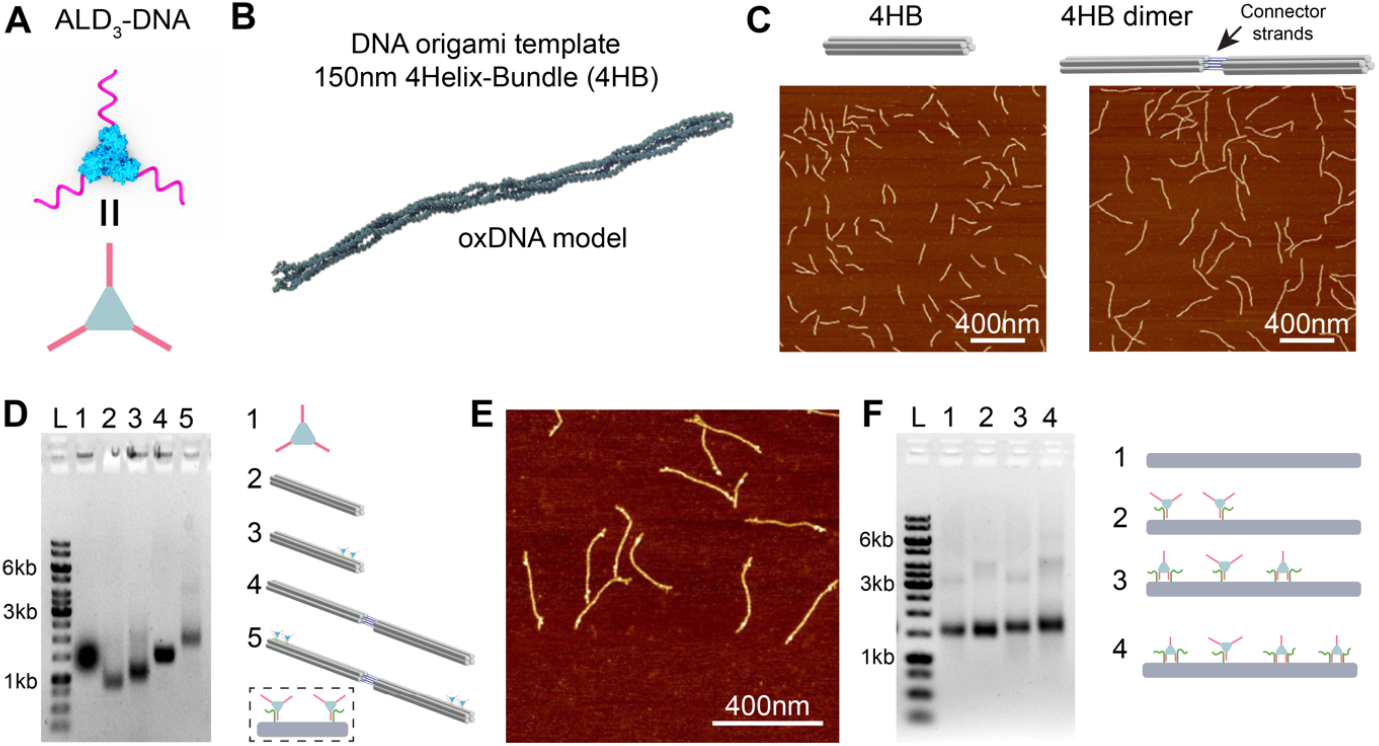
Synthesis of ald_3_-DNA and 4HB DNA origami template. (A) Schematic of ald_3_-DNA. The three DNA strands on the trimer are identical. (B) 4HB DNA origami. (C) AFM images of 4HB monomer and dimer. (D) Native agarose gel shows ald_3_-DNA binding to 4HB monomer and dimer. Lane L: 1kb DNA ladder. The numbered cartoons indicate the species in the corresponding well. (E) AFM image of purified 4HB dimer with two ald_3_-DNA on each origami. (F) Agarose gel of 4HB dimers with two, three, and four ald_3_-DNA. Lane L: 1kb DNA ladder. All schematic drawings show only a monomer of the DNA origami template. For simplicity, this is applied to the following figures as well.

The 6-nm ald_3_ trimer is very small compared to a typical DNA origami nanostructure, which can be hundreds of nanometers in one dimension. Furthermore, the 3 nm thickness is too thin to be easily visualized on origami structures that consist of multiple layers. To more readily observe the proteins tethered to the origami structure, we designed a relatively small 4-helix bundle (4HB) origami, using a 1800 nucleotide (nt) plasmid scaffold^42^ (**Figure 2B, Supplementary Figure S2**). The 4HB is 150 nm in length and only 5 nm in thickness. This size, and especially the modest cross section, makes it easier to see proteins arranged on the origami in Atomic Force Microscopy (AFM) images. To assemble the origami, the scaffold (50 nM) and 5 equiv of staples were mixed in 1×TE and 10mM MgCl_2_ buffer. The mixture was incubated at 95°C for 5 min, then annealed from 65°C to 2°C over 3.5 h. We synthesized both 4HB monomers and dimers for protein attachment. The latter was used in most of experiments, for reasons to be discussed below. The origami monomer and dimer were purified using 0.67% native agarose gels, and imaged by AFM (**Figure 2C**). DNA strands for the attachment of ald_3_-DNA were included in the assembly of 4HB origami. First, we tested attachment of two proteins to a 4HB monomer with one attachment strand, and the distance between two proteins is 40-bp along the 4HB (**Figure 2D**). After purification, 10 equiv per binding site of ald_3_-DNA was added to the 4HB. To ensure efficient protein binding, the mixture was allowed to incubate at 25°C for 48 h. Next, the sample was extracted from a 1% agarose gel to remove free ald_3_-DNA. Unfortunately, after attaching the ald_3_-DNA to the 4HB, the protein-origami conjugate has a mobility very similar to that of the free ald_3_-DNA, complicating the purification process (**Figure 2D**). To overcome this obstacle, we engineered 4HB dimers (300-nm total length) by connecting two identical nanostructures, tail-to-tail, via four sticky ends (**Supplementary Figure S2**). Because the 4HB dimer showed much slower mobility then free ald_3_-DNA, the dimer with proteins can be successfully separated from unbound proteins (**Figure 2D**). Further evidence for the separation was observed in the AFM image of the purified ald_3_-4HB dimer, which showed no proteins in the background (**Figure 2E**). For these reasons, 4HB dimers were used in all following experiments; however, for simplicity, all schematics only depict a single monomer. Attachment of two, three, or four ald_3_-DNA onto 4HB origami dimer with either one or two attachment DNA strands was successful, showing the robustness of this approach (**Figure 2F**).

We next explored the optimal length of “connector” DNA – the DNA duplex for connecting ald_3_-DNA building blocks. The connector consisted of two DNA strands that form a partially complementary duplex (whose length can be tuned), and two identical sticky ends complementary to the ssDNA handles on the protein. We reasoned that the middle domain length should not be too long (which would increase the difficulty of protein binding), but should also be greater than 15-bp to ensure thermal stability at room temperature. We were especially concerned that—since both ends of the connectors can bind to handles on the protein— an undesired *intramolecular* connection might result, forming a loop on a single ald_3_-DNA conjugate, and abrogating any further connections (**Figure 3A**). Our first test used three connectors with middle domain lengths of 15-bp, 25-bp, and 35-bp, which could bind with ald_3_-DNA modified with 21-nt handles, as reported in a previous study^39^. We mixed ald_3_-DNA and connectors in molar ratios of 1:1.5 incubated the samples at room temperature overnight, then analyzed them using an 8% native PAGE gel (**Figure 3B**). If the intermolecular connection dominates, aggregation will occur after mixing the ald_3_-DNA and connectors. In all cases, no aggregation was observed, and the higher intensity bands indicate one ald_3_-DNA binding with one or more connectors. Even though these bands suggested that the intermolecular connections partially occurred, the yield was not high enough for subsequent experiments. Shortening the connectors will increase its rigidity and therefore can potentially reduce its tendency of forming an intramolecular loop. To decrease the total length of the connector, we modified the ssDNA attached to ald_3_ from 21 to 15 nt, while keeping the middle section at either 15-bp or 21-bp. Ald_3_-DNA and connectors were mixed and incubated. Aggregation was observed in all ratios with the 15-bp connector. Also, protein multimers bound with the connector increased (**Figure 3C**). Although we still observed some individual protein-DNA conjugates with connectors, we hypothesized that these correspond to the ald_3_-DNA binding a single connector sticky-end, without forming a loop. Thus, all subsequent experiments used the 15-bp connector to create protein multimer nano-shapes. The product of ald_3_-DNA connected with DNA connector without DNA origami template was verified by AFM imaging (**Supplementary Figure S3**). As expected, protein assemblies of various sizes and geometries were observed in the images.

**Figure 3:**
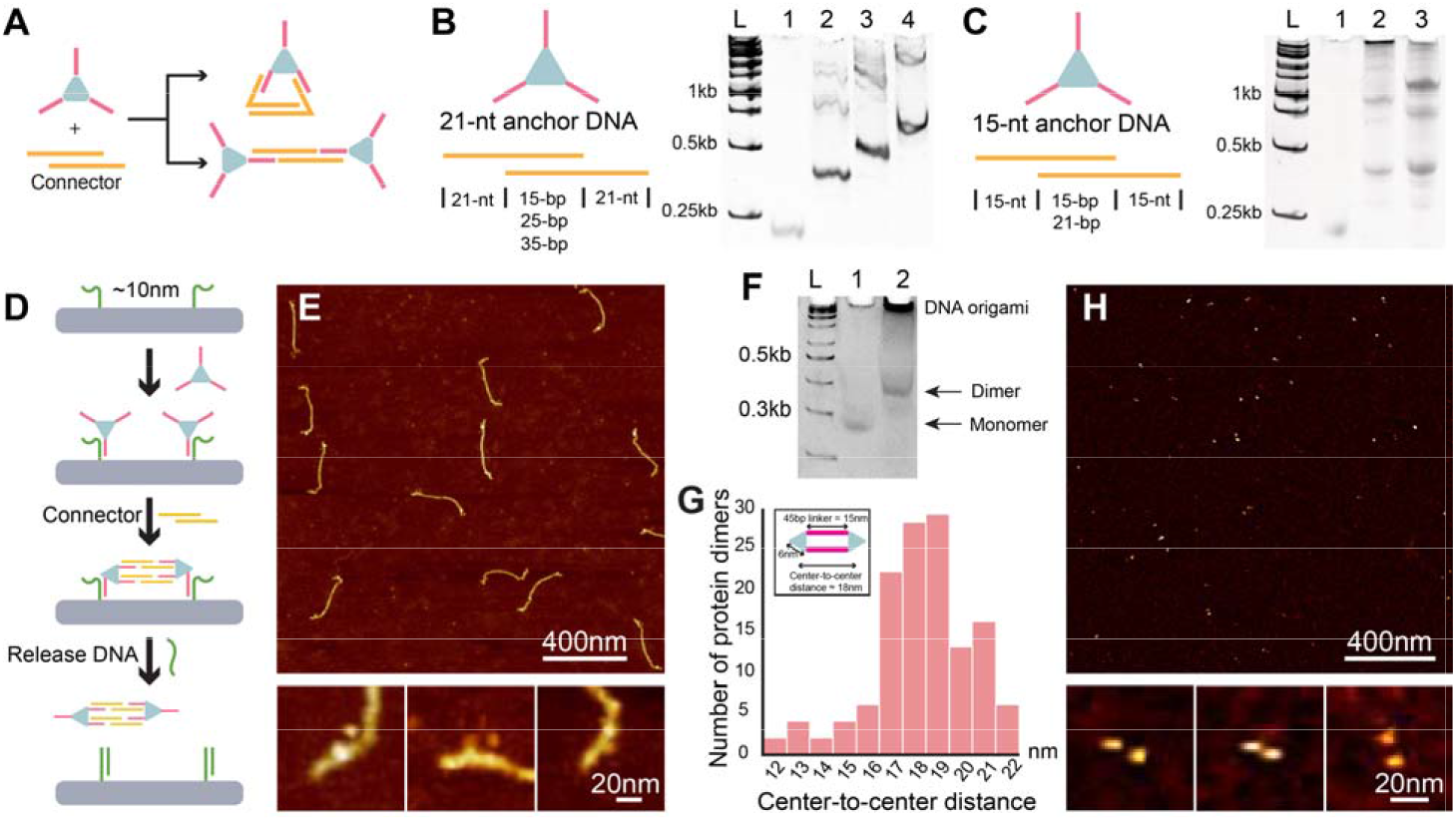
Optimization of connector DNA and formulation of ald_3_-DNA dimer. (A) Connector DNA can form intra-(top) or inter-molecular (bottom) connections. The connector needs to be optimized to promote the desired intermolecular connection. (B) Optimization of the middle duplex region of the connector by using an ald_3_-DNA with a 21nt long ssDNA. In native PAGE, lane L: 1kb DNA ladder; lane 1: ald_3_-DNA; lane 2: ald_3_-DNA with short connector DNA (15bp duplex in the middle); lane 3: ald_3_-DNA with medium connector DNA (25bp duplex in the middle); lane 4: ald_3_-DNA with long connector DNA (35bp duplex in the middle). The leading bands in lanes 2, 3, and 4 are monomers that formed intramolecular connection. (C) Optimization of the middle duplex region of connector by usin an ald_3_-DNA with a 15nt long ssDNA. lane L: 1kb ladder; lane 1: ald_3_-DNA; lane 2: ald_3_-DNA with short connector DNA (15bp duplex in the middle); 3: ald_3_-DNA with long connector DNA (21bp duplex in the middle). The aggregate indicates formation of intermolecular connection. (D) Schematic of molecular operations for dimer assembly and release on DNA template. (E) AFM images of ald_3_-DNA dimers assembled on 4HB DNA origami. (F) Native PAGE gel of ald_3_-DNA dimer after being released from 4HB. Lane L: 100bp ladder; lane 1: ald_3_-DNA; lane 2: ald_3_-DNA dimer after release. The aggregate in the loading well in lane 2 is the DNA origami scaffold, which is too large to enter the gel (G) Histogram of center-to-center distance for ald_3_-DNA dimers. Inset shows the estimated distance based on the dimer structure. (G) AFM images of ald_3_-DNA dimers.

After optimization of DNA origami template and DNA connectors, we next probed formation of a protein dimer on the 4HB template. Each attachment strand for immobilizing the ald_3_-DNA to the origami is 35 nt long (15 nt complementary to the handle on the protein, and another free 20 nt) in length. While the ends closest to the origami surface bind to the ssDNA on ald_3_-DNA, a single-stranded domain was left unpaired on the 3’ end to enable displacement of the protein after linking the proteins with the connector strand (**Figure 3D**). Successful attachment of two proteins to 4HB dimer was verified with AFM imaging after gel purification (**Figure 3E**). The ald_3_-DNA dimer can be released from the 4HB via addition of a release DNA strand, which includes a 20-nt complementary sequence to the attachment strand toeholds and a 10-nt domain hybridizing with the protein binding domain. After the final products are released from the origami, leftover connectors in solution can bind to the released protein multimer and further cause aggregation. To avoid this issue, a 15-nt “blocker” strand was added to deactivate the extra connector before the last releasing step (**Supplementary Figure S4**, See SI for dimer assembly in detail). Regarding the blocker DNA, PAGE analysis showed little difference between addition vs. omission of the blocker strand. However, after recovering the product from the gel and analyzing the sample by AFM, extensive aggregation was observed for the ald_3_-DNA dimer without the blocker strand. This observation confirmed our hypothesis that the leftover connector can hybridize with free handles on the released protein nanostructures, and blocker strands were used in subsequent experiments. After being released from 4HB template, ald_3_-DNA dimers were purified by 8% native PAGE (**Figure 3F**), and visualized in AFM images after purification from the gel (**Figure 3G and 3H, Supplementary Figure S5**). The average distance between two ald_3_ is 17.3 ± 0.2 nm (SD, N = 132), close to the calculated distance of ∼18nm. Although the assembly of dimers on 4HB appeared to be very good, the yields of free dimers in PAGE could not be quantified, because aggregation of DNA origami in the loading wells. As a result, we calculated the percentage of complete dimer (proteins in dimer formation/total number of proteins) after PAGE purification and obtained a yield of 84% based AFM image analysis. This method was used for estimating the yields of other multimer formations as well.

Assembly of the ald_3_-DNA dimer was realized by anchoring each protein building block on 4HB with a single attachment DNA strand. This approach worked well for dimer formation because it allowed the maximum flexibility for connection (i.e., the protein can move more freely to accommodate the connector DNA) and there is only one possible way to link the ald_3_-DNA building blocks. For more complex assemblies, such as the linear ald_3_-DNA trimer, the single-attachment method can lead to dimer formation and prevent subsequent linking into a trimer (**Figure 4A, Supplementary Figure S6**). To avoid this issue, we used a combination of single attachment and double attachment of ald_3_-DNA to the 4HB template to eliminate unwanted side reactions (**Figure 4A**). After assembly and connection of liner trimer, the product can be freed from DNA template by adding a release DNA strand (**Figure 4B**). Another issue, however, arose from this new scheme: the double attachment of ald_3_-DNA is expected to limit the freedom of protein units. Therefore, it is important to carefully design and optimize the inter-protein distances to achieve efficient assembly. The distance between proteins along the 4HB should be large enough to prevent the “edge” proteins from binding to the handles for the “middle” ald_3_-DNA. Conversely, the distance should be small enough for the connector to link the building blocks. Two distances (40 bp and 32 bp) between attachment sites were tested (**Figure 4C**). After a 48-hour incubation to allow all three protein-DNA conjugates to bind, successful binding of three ald_3_-DNA on the 4HB for both designs were confirmed with agarose gel electrophoresis (**Supplementary Figure S7**). Nonetheless, after adding connector DNA and release DNA, the final yields of liner trimers for the two designs are drastically different, although both the 40-bp and 32-bp inter-protein distances should be shorter than the connectors. The 32-bp distance generated a clear trimer ald_3_-DNA band in native PAGE, while the 40-bp sample showed a barely visible trimer band. As a result, we selected the more suitable 32 bp as the final distance between two proteins.

**Figure 4:**
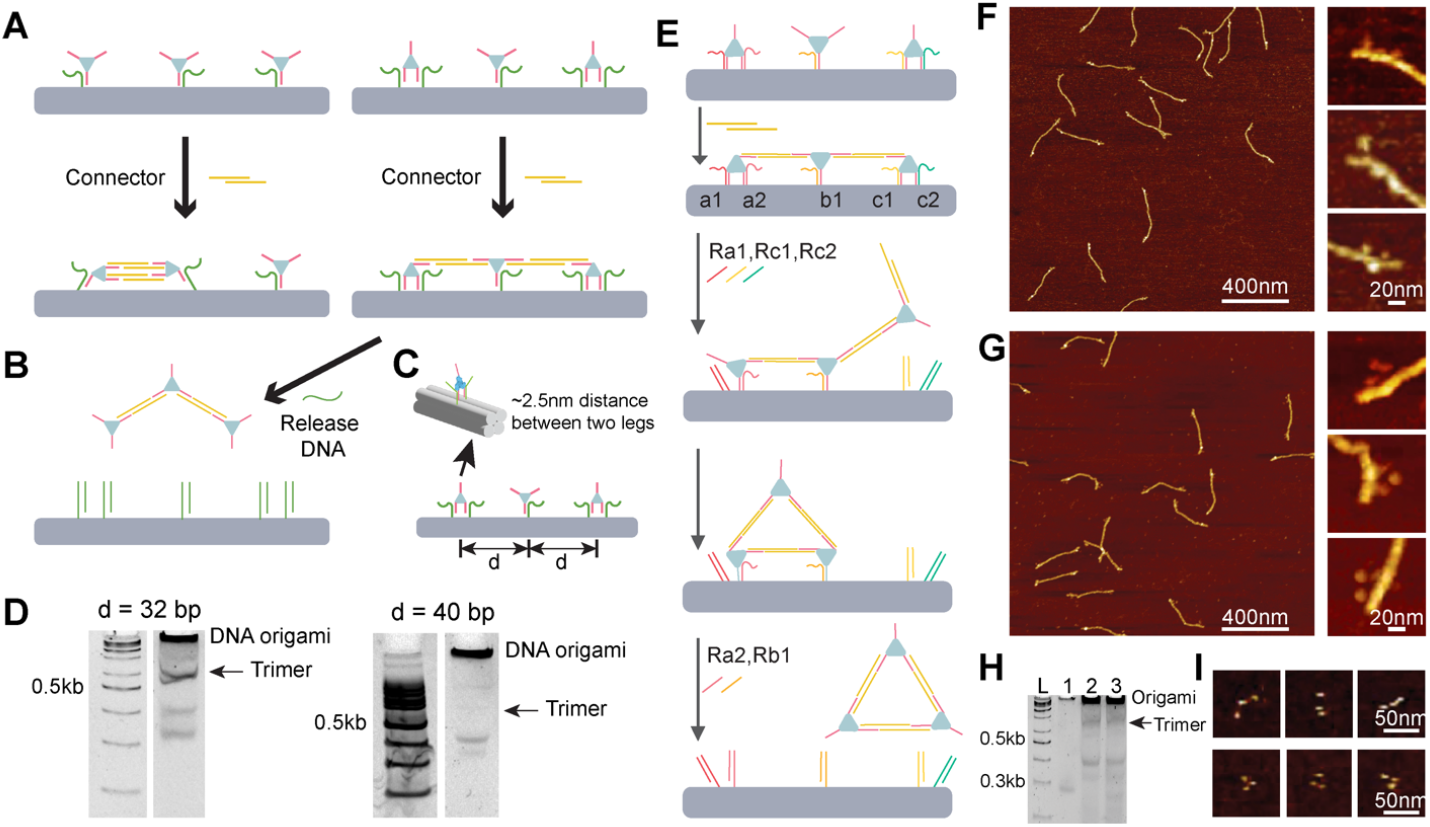
Assembly of ald_3_-DNAtrimers. (A) Design scheme for attaching three proteins on DNA template and connecting them to form a linear trimer. (B) Linear trimer ald_3_-DNA can be released from DNA template. (C) Optimization of inter-protein distance for more effective connection. (D) PAGE analysis of released products generated from two 4HB templates (d = 32bp and 40 bp). Arrows in PAGE indicate the position of ald_3_-DNA trimer. The 32bp distance showed significantly higher yield. (E) Molecular operation scheme for forming a triangle trimer. (F) and (G) AFM images of linear trimers and triangle trimers on 4HB DNA origami. (H) PAGE analysis of free linear trimers and triangular timers after being released from DNA origami. Lane L: 100 bp DNA ladder; Lane 1: ald_3_-DNA; Lane 2: linear trimer; Lane 3: triangle trimer. Arrow indicates the position of ald_3_-DNA trimers. (I) Zoomed-in AFM images of linear trimers and triangle trimers.

We next moved to assembling a triangular ald_3_-DNA shape. Unlike the linear trimer, which was connected in a single-step reaction, constructing an ald_3_-DNA triangle requires a more complex set of molecular operations. Taking advantage of the programmability of DNA origami template, we used a sequential, multi-step reaction that frees up specific DNA strands at different stages to allow intended connections (**Figure 4E**). To realize the stepwise release necessary for this structure, it is necessary to have different sequences for the attachment vs. releasing strands. Since the handles on the protein must be identical, this orthogonality is encoded in the single-stranded toehold domains of the docking strands on the origami, allowing specific legs of the ald_3_-DNA to be released at a desired time. After forming a linear trimer on the 4HB, we released one leg of the leftmost protein (***a1***) and *both* legs of the rightmost protein (***c1, c2***). ***Ald c*** is thus free to swing around and bind to ***ald a*** using a connector strand in solution. As a result, a triangle forms on the 4HB and can be liberated using subsequent releasing strands. The only difference between the synthesis of the linear trimer and the triangle is the addition of releasing strands ***Ra1, Rc1***, *and* ***Rc2*** for the formation of triangle; we note that an excess of free connector DNA has been in solution, enabling connection of the ald_3_-DNA as soon as they are freed by the releasing strands). We can then add blocker strands followed by the releasing strands, ***Ra2*** and ***Rb1***, to liberate the fully formed triangle (See SI for reaction information in detail). AFM images confirmed both linear trimer and triangle trimer assembled on 4HB template (**Figure 4F and 4G**). After release, the free trimer products were purified with PAGE (**Figure 4H**), and imaged with AFM (**Figure 4I**). After purification of free trimer products from PAGE, we estimated the percentages of complete trimers as 78% and 45% for the linear timer and triangle timer, respectively (**Supplementary Figures S8 and S9**).

The formation of a linear trimer vs. triangle nanostructure demonstrated the feasibility of using an origami to sequentially dictate the nanostructure formed, from a common set of identical building blocks. We next asked whether we could extend this method to assemblies of four ald_3_-DNA. Unlike the ald_3_-DNA trimer, which has a linear isomer and a non-linear isomer, a tetramer of protein building blocks allows for a linear isomer and two non-linear tetrameric isomers: a square and a triangle with a single protein linked to one corner (which for simplicity we term the “Y-shape”). Both structures, however, first require assembly of a linear tetramer on the 4HB. To bind four proteins together into a linear tetramer (in the order ***abcd***) requires one free DNA leg from building blocks ***a*** and ***d*** and two legs from ***b*** and ***c***. However, ***b*** and ***c*** cannot have two free legs from the outset, or else they may form a double connection to one another, thereby preventing the formation of linear tetramers (**Figure 5A**). When arranging four ald_3_-DNA on the 4HB, ***a, c***, and ***d*** have two legs bound and ***b*** has only one leg bound. Two connections form after adding connectors. In one situation, ***a, b***, and ***c*** are connected as a linear trimer, while ***d*** remains unconnected. In another case, ***a*** and ***b***, as well as ***c*** and ***d*** are linked as two dimers, and one leg of ***b*** is not connected to another protein. For both situations, releasing one leg of ***c*** can help it connect with the remaining legs and form a linear tetramer. At this step, adding all other six releasing strands would lead to a linear ald_3_-DNA tetramer. However, instead of releasing the linear trimer, two assembly protocols could be implemented to make either the square or Y-shape (**Figure 5B**). With the linear tetramer on the 4HB, releasing ***a1, a2, d1, d2***, then linking ***a*** and ***d*** will result in a square. Releasing ***b1*** and ***c1*** at this point liberates the square into solution. Alternatively, releasing ***b1, d1***, and ***d2*** first, then connecting ***b*** connects with ***d***, and releasing ***a1, a2, c1*** will result in the Y-shape. It is more difficult to observe all four ald_3_-DNA components on the origami (compared with only two or three), and the proteins are obscured and hard to distinguish given their small size and close proximity. Nonetheless, we could identify structures that clearly showed four protein building blocks on the end of 4HB (**Figure 5C**). In AFM images of the square and Y-shape, we can often directly observe the formation of the corresponding geometries formed by ald_3_-DNA (**Figure 5C**). The native PAGE gel showed that all three tetramers have similar mobility (**Figure 5D**), and AFM imaging of the released, purified tetramers clearly showed expected formations of the ald_3_-DNA building blocks (**Figure 5E**). The percentages of complete tetramer were estimated to be 45%, 35%, and 33% for the linear tetramer, the square tetramer, the Y-shaped tetramer, respectively (**Supplementary Figures S10 to S12**). Add DNA sequences used in this work can be fold in Supplementary Information.

**Figure 5:**
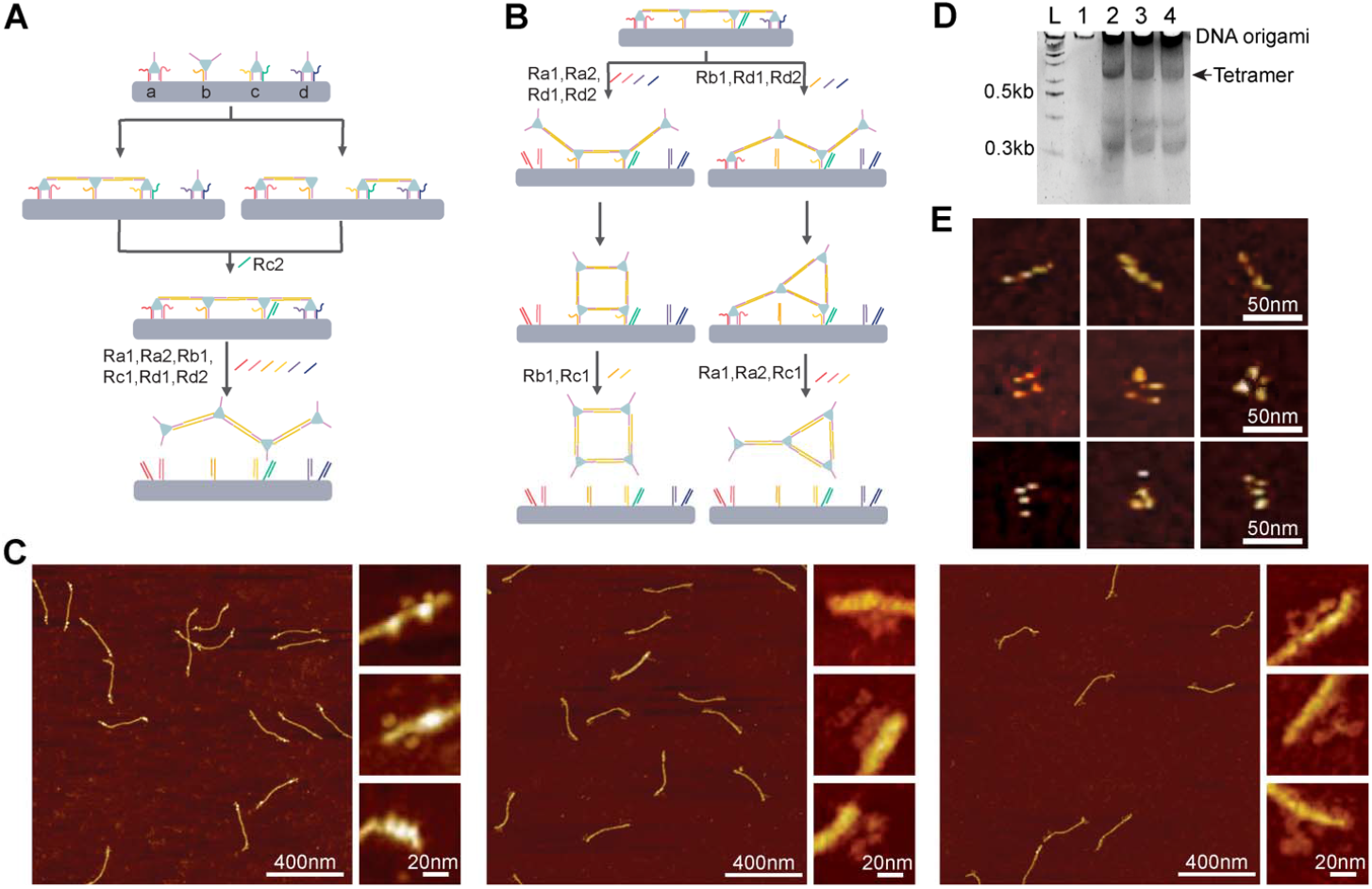
Assembly of ald_3_-DNA tetramers. (A) Scheme for linear tetramer assembly. (B) Scheme for the assembly of square tetramer and Y-shaped tetramer. (C) From left to right, AFM images of linear tetramer, square tetramer, and Y-shaped tetramer on 4HB DNA origami. (D) PAGE analysis of free linear tetramer, square tetramer, and Y-shaped tetramer after being released from DNA origami. Lane L: 100 bp DNA ladder; Lane 1: ald_3_-DNA; Lane 2: linear tetramer; Lane 3: square tetramer; Lane 4: Y-shaped tetramer. Arrows indicate the position of ald_3_-DNA tetramers. (E) From top to bottom, zoomed-in AFM images of linear tetramer, square tetramer, and Y-shaped tetramer.

## Conclusions

To summarize, we have developed an approach that uses DNA origami as a platform, with ssDNA-modified oligomeric proteins as a common building block, and performs sequential strand displacement and re-connection operations to yield isomeric protein oligomers. Only one type of protein-ssDNA conjugate was used here, and all three DNA handles bound to the protein trimer share the same sequence. The protein-ssDNA building block alone cannot produce designable, precise assembly. Nevertheless, we successfully linked protein-DNA conjugates to create dimers, two trimeric isomers, and three tetrameric isomers. The DNA origami template not only spatially segregates the protein building blocks to prevent incorrect (or premature) association, but the handles for attaching the protein to the origami have unique single-stranded regions that enable specific release of only the desired handles at desired times. Thus, the final structures obtained are “encoded” in: (1) the number of docking sites on the origami; (2) the single-stranded regions on the origami docking strands; and (3) the order of displacement and connection strands added.

This work breaks from previous reports of protein-DNA conjugates, which yielded polydisperse assemblies like nanofibers or 3D crystals, to generate unique numbers and connectivity of protein oligomers through programmatic DNA-DNA connections. A number of additional, more complex assemblies can be envisioned by using two populations of ald_3_-DNA, each with different DNA handles (which would allow for unique positioning on the origami). Taking the tetramer as an example, if the legs of ald_3_-DNA ***c*** have a different sequence from the ***a, b***, and ***d*** building blocks, two legs can be free for connection, and a linear tetramer can be formed in only one connection step (**Supplementary Figure S13**). Also, in this work we demonstrated protein complex attachment a linear, pseudo-1D origami scaffold. Much greater topological opportunities are available by moving to a more complicated DNA nanostructure. Furthermore, ald_3_ is not the only protein that can be conjugated to ssDNA and assemble as homomultimeric protein complexes later. A range of other protein oligomers with varying symmetries could be used; alternatively, developing methods for modifying a single oligomer with two or more ssDNA sequences would offer further complexity in the nanostructures formed. Our approach also in principle allows for the generation of hetero-multimeric protein complexes if more than one *type* of protein oligomer is used (e.g., combining ald_3_-DNA with a tetrameric aldolase^43^ modified with different DNA handles). Indeed, ultimately a series of Lego-like protein building blocks with addressable handles could be hierarchically connected through supramolecular linkages, much the way that various hybridization states of carbon (sp, sp^2^, sp^3^) are linked by organic synthesis into molecules with complex presentation of functionality in 3D space.

The protein complexes developed in this work do not have a specific function, but rather serve only as a structural nanomaterial. However, these materials can find broad applicability by incorporating functional proteins, either by fusing them genetically to the oligomeric building blocks, or by chemical or enzymatic attachment (e.g., via SpyCatcher/SpyTag^44^) after the fact. Current approaches for constructing multi-protein complexes requires engineering, or genetically grafting, a protein-protein interface, which can be difficult for making complex shapes. Our approach breaks this restriction, and allows a range of nanostructures to be constructed from a simple set of common building blocks, thanks to the programmability of DNA linkers and the spatial restrictions imposed by the origami scaffold. We ultimately envision applications such as multienzyme complexes performing a composite biocatalytic reaction, nanopores in biological membranes, synthetic antibodies of tunable shape and size, drug delivery vehicles, synthetic multivalent protein vaccines, and other biomolecular nanostructures.

## Acknowledgements

This work was supported by the National Science Foundation grant CCF-2227399 to Y.K., and by the National Science Foundation DMR-BMAT CAREER award 1753387 and the National Institute of General Medical Sciences of the National Institutes of Health grant DP2GM132931 to N.S. The content is solely the responsibility of the authors and does not necessarily represent the official views of the National Institutes of Health.

