## Supplementary information for "DNA Nanostructure-Guided Assembly of Proteins into Programmable Shapes"

**Materials and methods**

**Materials.** The single-stranded DNA scaffold (pscaf1800) was extracted from a plasmid. Chemically synthesized DNA short strands were purchased from Integrated DNA Technologies (www.idtdna.com) and were used without further purification. All other reagents were purchased from Sigma-Aldrich.

**DNA origami design and folding.** The 4HB DNA origami was designed with the software caDNAno (http://cadnano.org/). For DNA Origami folding, 10 nM scaffold together with a tenfold excess of each staple strand was mixed in 1×TE (10 mM Tris, 1 mM EDTA; pH 8.0) buffer with 10 mM MgCl_2_. In the annealing process the folding mixture was heated at 80°C and slowly cooled from 65°Cto 25°C at a rate of -1°C/5min. Afterwards, the folded DNA origami was purified using agarose gel electrophoresis.

**Agarose gel electrophoresis.** DNA origami samples were subjected to agarose gel electrophoresis at 75V for 2-3 hours in an ice water bath. Gels were prepared with 0.5× TBE buffer containing 10 mM MgCl_2_ and 0.005% (v/v) Ethidium Bromide.

**Protocol for dimer assembly.** 5 equiv of connector per pair of ald_3_-DNA building blocks (i.e., per dimer) was added to the sample and allowed to incubate at 25°C overnight. Next, 4 equiv blocker per connector (corresponding to 20 equiv per dimer) was added, followed by incubation for 1 h. Finally, 5 equiv of releasing strands per attachment strand (so 10 equiv per dimer) was added for 3 h to release the dimer.

**Protocol for trimer assembly.** The protocol for formation of the linear trimer is the same as for the dimer. To form the triangle trimer, the linear trimer was synthesized first, then 5 equiv of releasing strands **Ra1, Rc1,** and **Rc2** was added, and the solution was incubated for at least 16h for this release and re-connection process. AFM images demonstrate that the ald_3_-DNA triangles formed in the expected locations (**Figure 5F**). We then added blocker strands followed by the releasing strands, **Ra2** and **Rb1**, to release the fully formed triangle.

**Protocol for tetramer assembly.** 10 equiv of connector was added to the sample and incubated at 25 °C overnight. We then added 5 equiv of releasing strands **Rc2,** and incubated for 16h to enable linear tetramer formation. To release the linear tetramer, we added 40 equiv blocker and incubated the sample for 1h, then added 5 equiv of releasing strands **Ra1, Ra2, Rb1, Rc1, Rd1** and **Rd2** for 3h. To form the square and Y-shape, we added (to the linear tetramer still bound to the origami) **Ra1, Ra2, Rd1, Rd2** for the square and ***R*b1, Rd1, Rd2** for the Y-shape, followed by another 16 h incubation. To release the square, we added strands **Rb1** and **Rc1**, while the Y-shape required strands **Ra1, Ra2** and **Rc1**; all strands were incubated for 3h, following an incubation for 1 h with 40 equiv of blocker strands.

**Native PAGE gel electrophoresis.** DNA oligonucleotides and protein samples were subjected to 8% native PAGE gel electrophoresis at 80V for 2-3 hours at room temperature. Gels were prepared with 0.5 ×TBE buffer containing 10 mM MgCl_2_, and stained with 1×Sybr Gold after gel running.

**Extraction of protein products from PAGE**. To recover a product, we cut the band and crush it with a small pestle at room temperature. For each band, 20 µL of elution buffer (500 mM Ammonium Acetate, 10 mM Magnesium Acetate, 2 mM EDTA) was added to the crushed gel to solubilize the protein products.

**AFM imaging.** To visualize samples by AFM, 2 μL of the solution was deposited onto freshly cleaved mica. The sample area was then filled with 80 μL1×TE buffer with 10 mM MgCl_2_. The samples were imaged on a Multimode VIII system (Bruker) in liquid using commercial tips (SNL-10, Bruker).

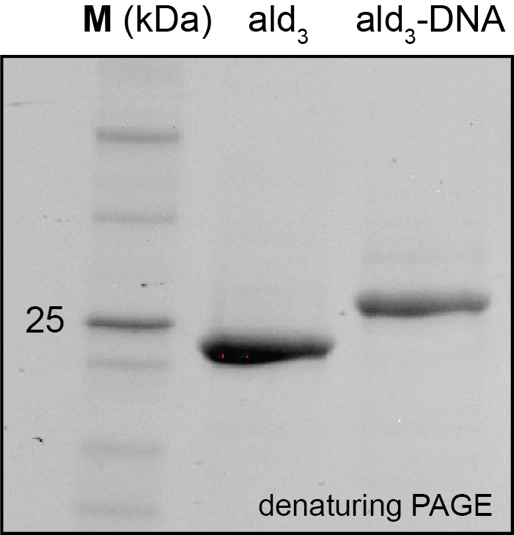

**Figure S1: Denaturing PAGE analysis of the protein-DNA conjugate.** The ald3-DNA conjugate shows a complete band shift from the unmodified protein, confirming that all three monomers of the trimer have been successfully conjugated to DNA."

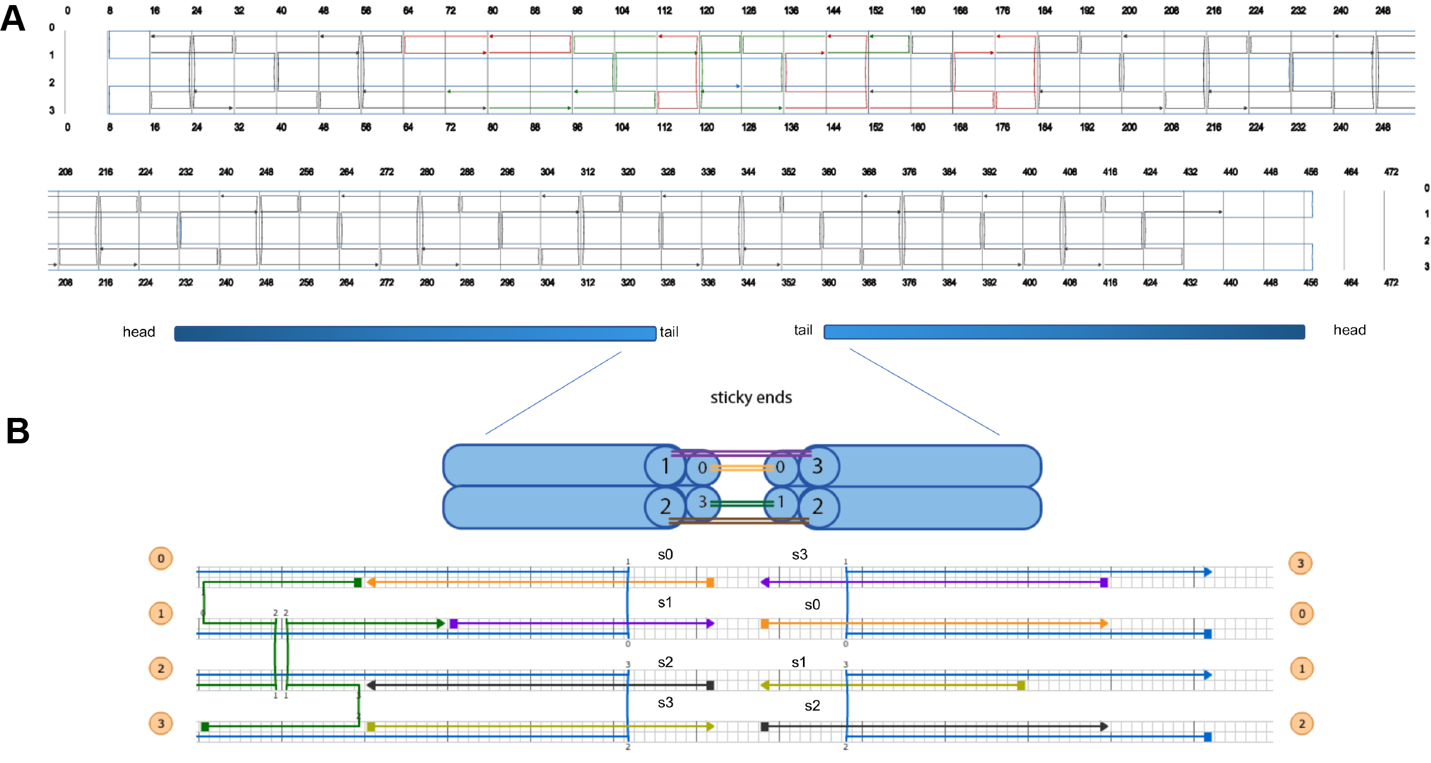

**Figure S2: 4HB origami and sticky ends design.** (A) CaDNAno design diagram of the 4HB origami. The red strands are attachment points, which can be modified. (B) Four sticky-end strands at the tail of 4HB. Sticky ends are 10nt long. S1 is complementary to s3, s0 and s2 are palindromic (self-complementary) sequences.

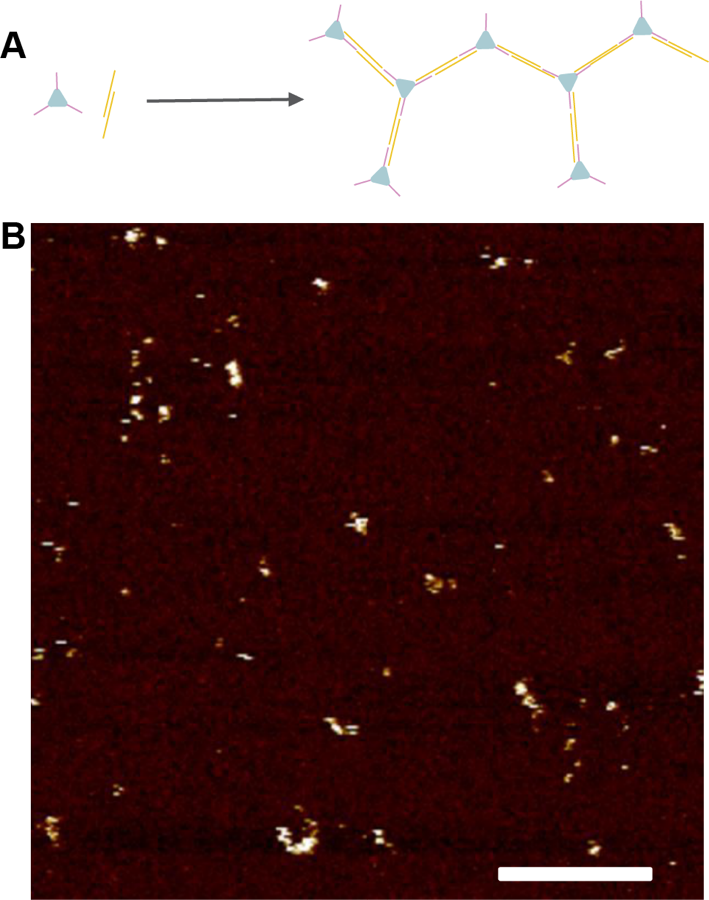

**Figure S3: Connecting ald_3_-DNA with connector DNA strands in absence of a DNA origami template.** (A) Illustration of ald_3_-DNA + connector random connection. (B) AFM image of ald_3_-DNA + 1.5eq connectors connected after 2h. Only ill-defined, heterogeneous aggregates are observed. Scale bar: 200nm.

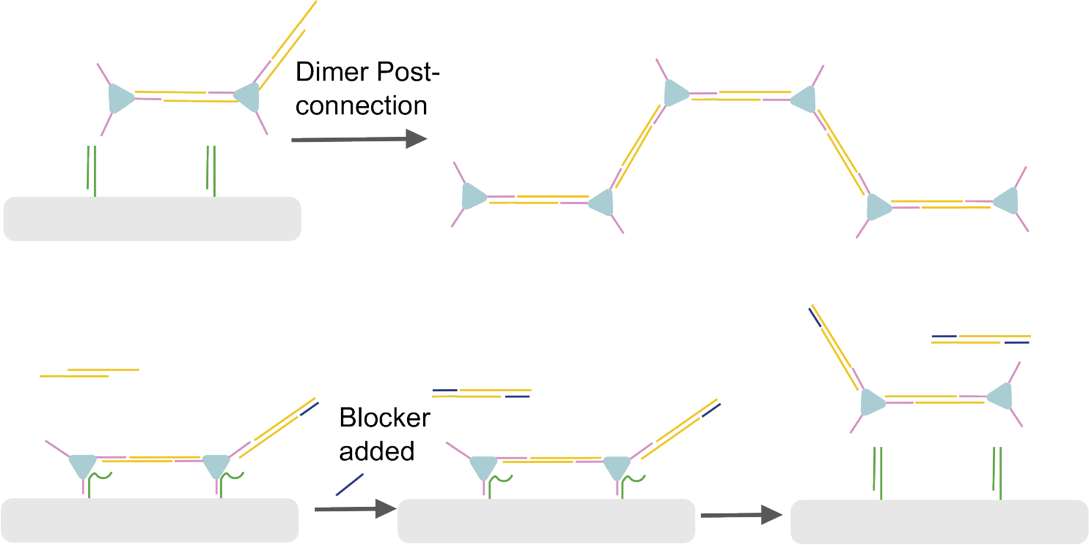

**Figure S4: Deactivation of the extra connector with blocker DNA.** Addition of the blocker strands prevents further spurious association of the dimer after connection (top), and ensures that only the desired dimer is isolated (bottom).

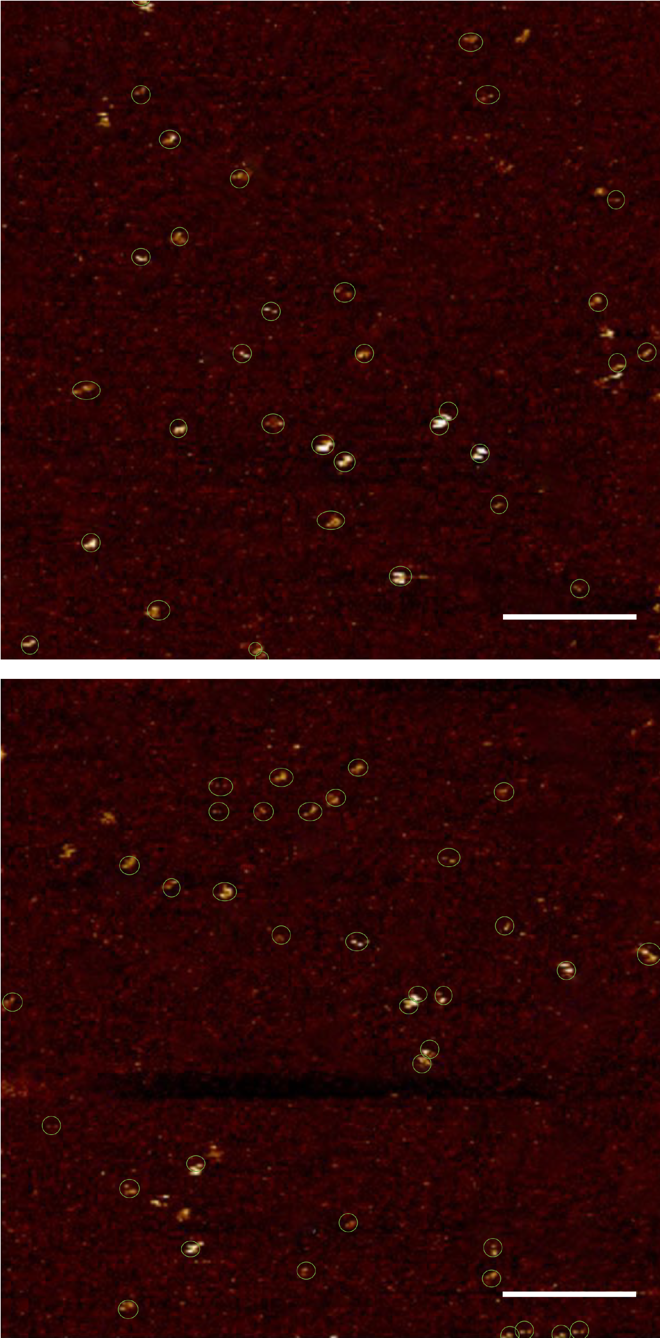

**Figure S5: Wide-field AFM images of released ald_3_-DNA dimers.** Ald_3_-DNA dimers are marked with green circles. Scale bars: 200nm.

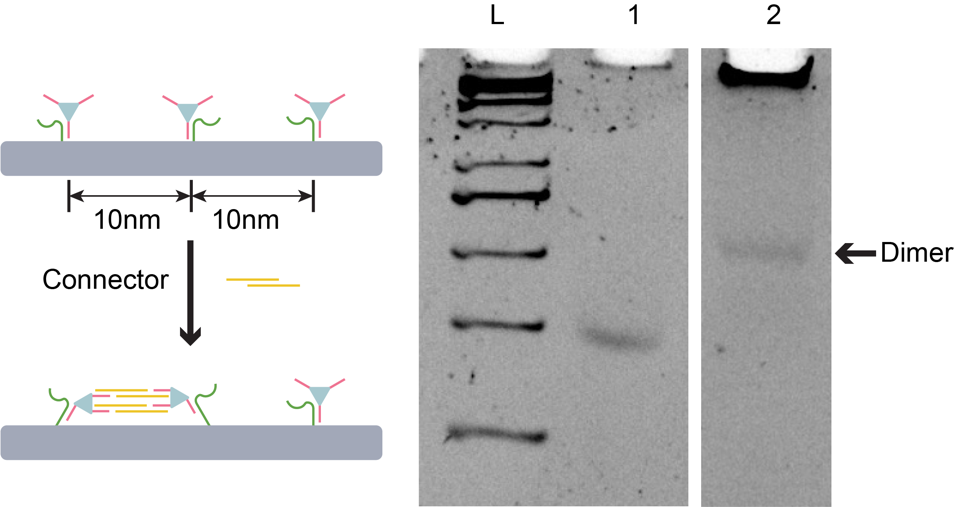

**Figure S6: Connecting three ald_3_-DNA units to 4HB with a single attachment produced only dimer products.** Left: Schematic of the assembly. Right: PAGE gel of products after releasing from 4HB. Lane L: 100bp DNA ladder. Lane 1: Ald3-DNA. Lane 2: Product. Only dimer was observed. As a result, the leftmost and rightmost ald_3_-DNA were attached to the origami at two points to avoid this dimer formation.

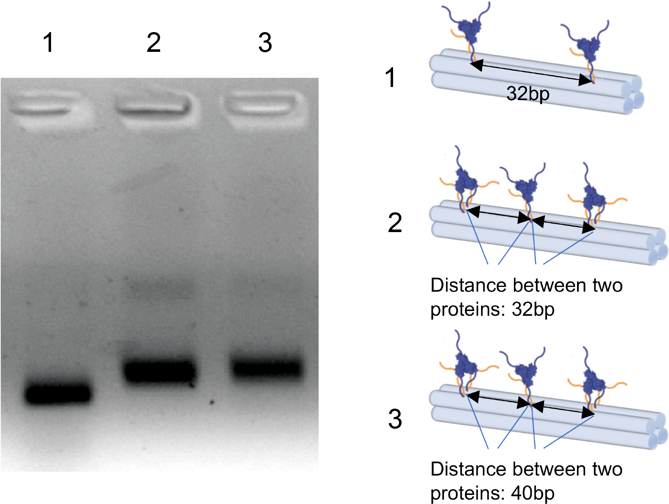

**Figure S7: Agarose gel analysis of assembly of three ald_3_-DNA on the 4HB template.** Sample 2 and sample 3 both showed slower mobility than control sample 1, which contains two ald_3_-DNA, suggesting most structures of sample 2 and sample 3 contain three ald_3_-DNA.

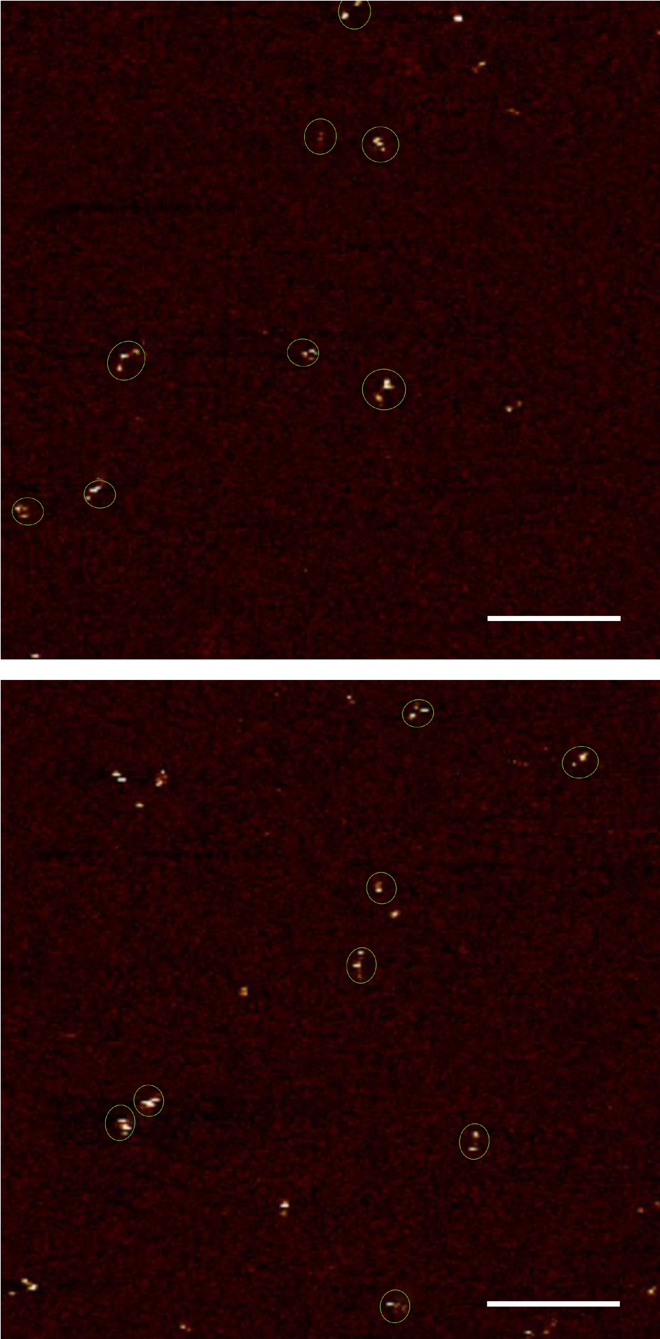

**Figure S8: Wide-field AFM images of recovered linear trimer.** Linear trimers are in green circles. Scale bars: 200nm

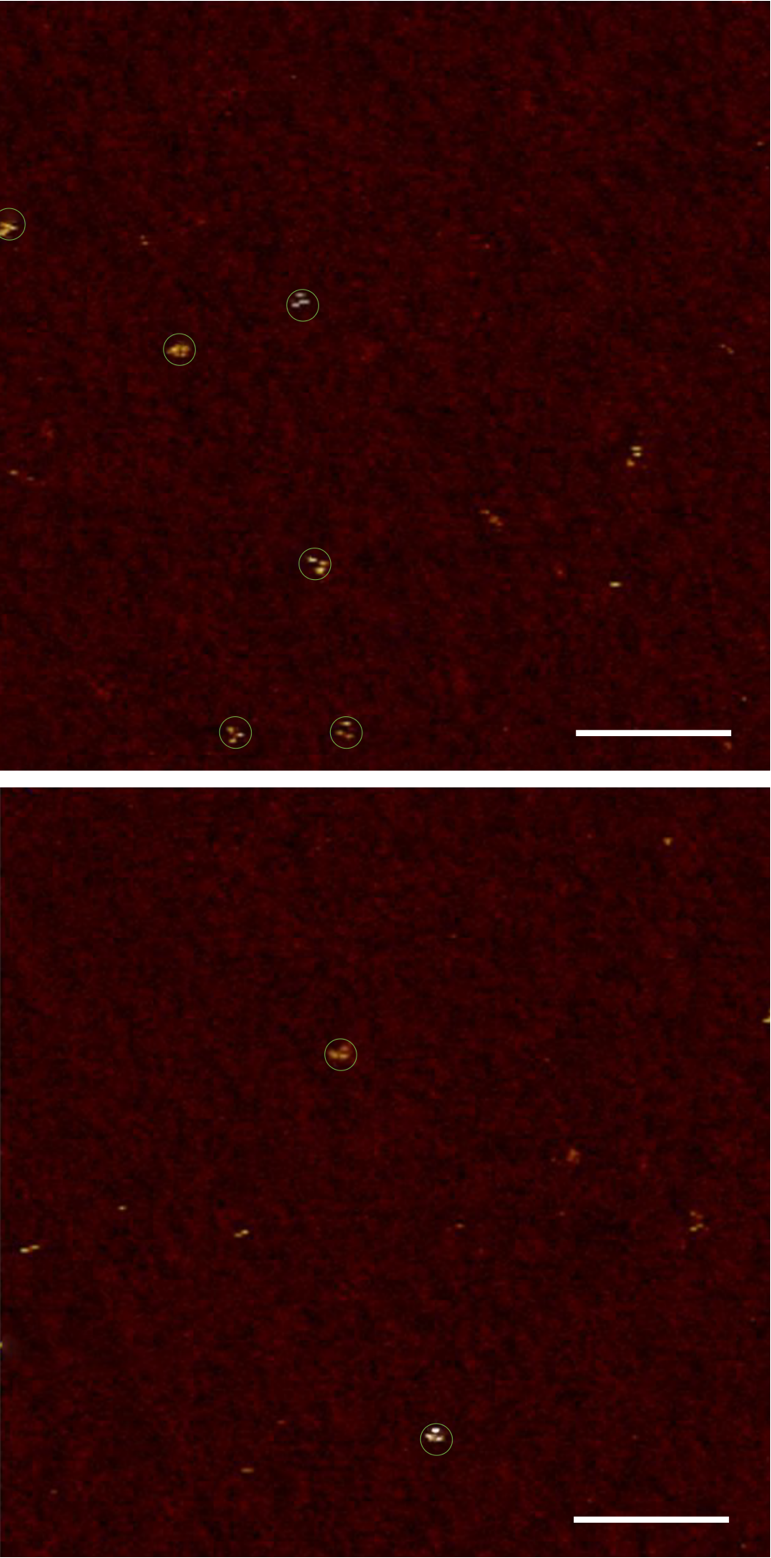

**Figure S9: Wide-field AFM images of recovered triangle trimers.** Triangle trimers are in green circles. Scale bars: 200nm.

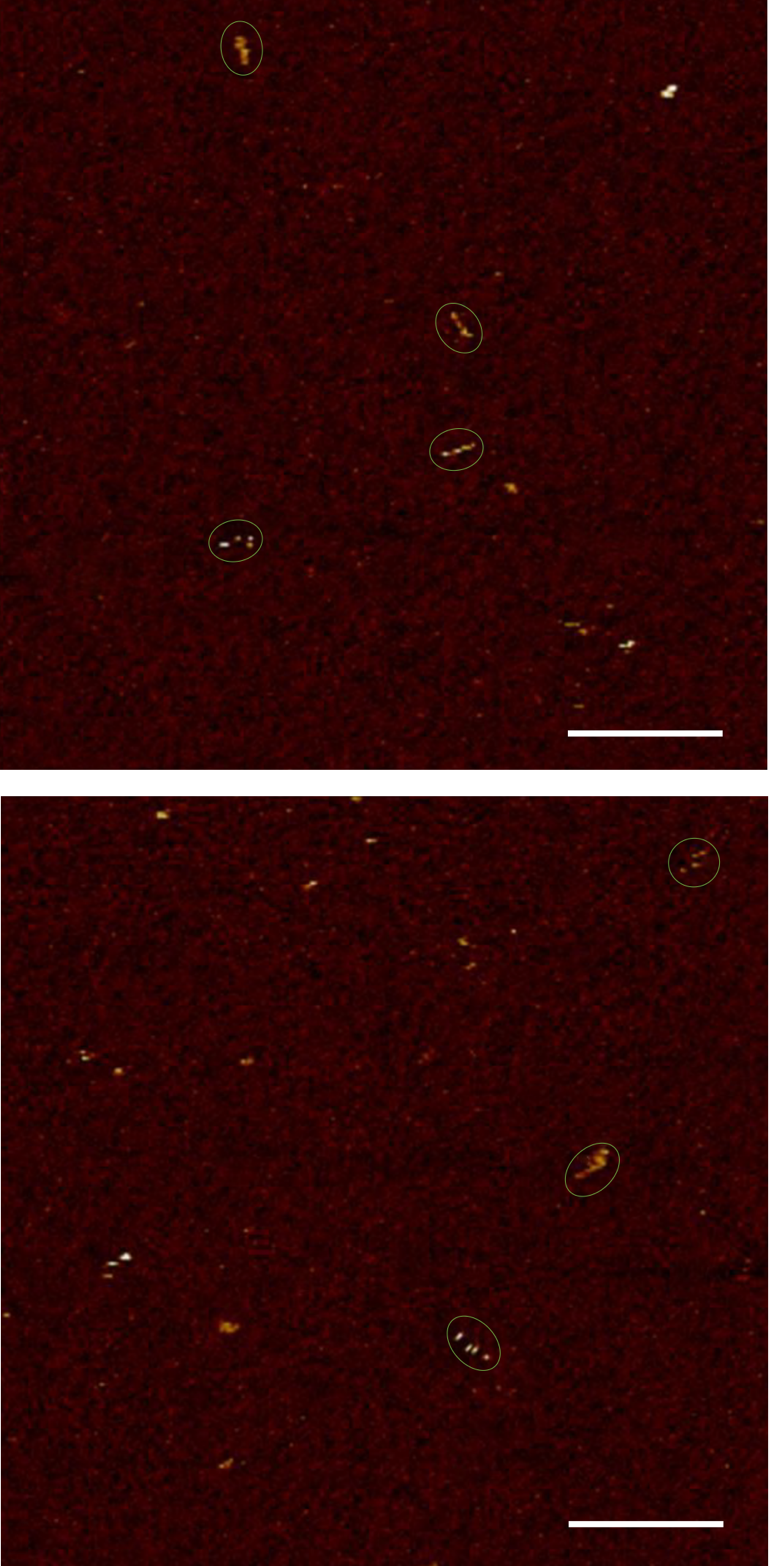

**Figure S10: Wide-field AFM images of recovered linear tetramer.** Linear tetramers are in green circles. Scale bars: 200nm.

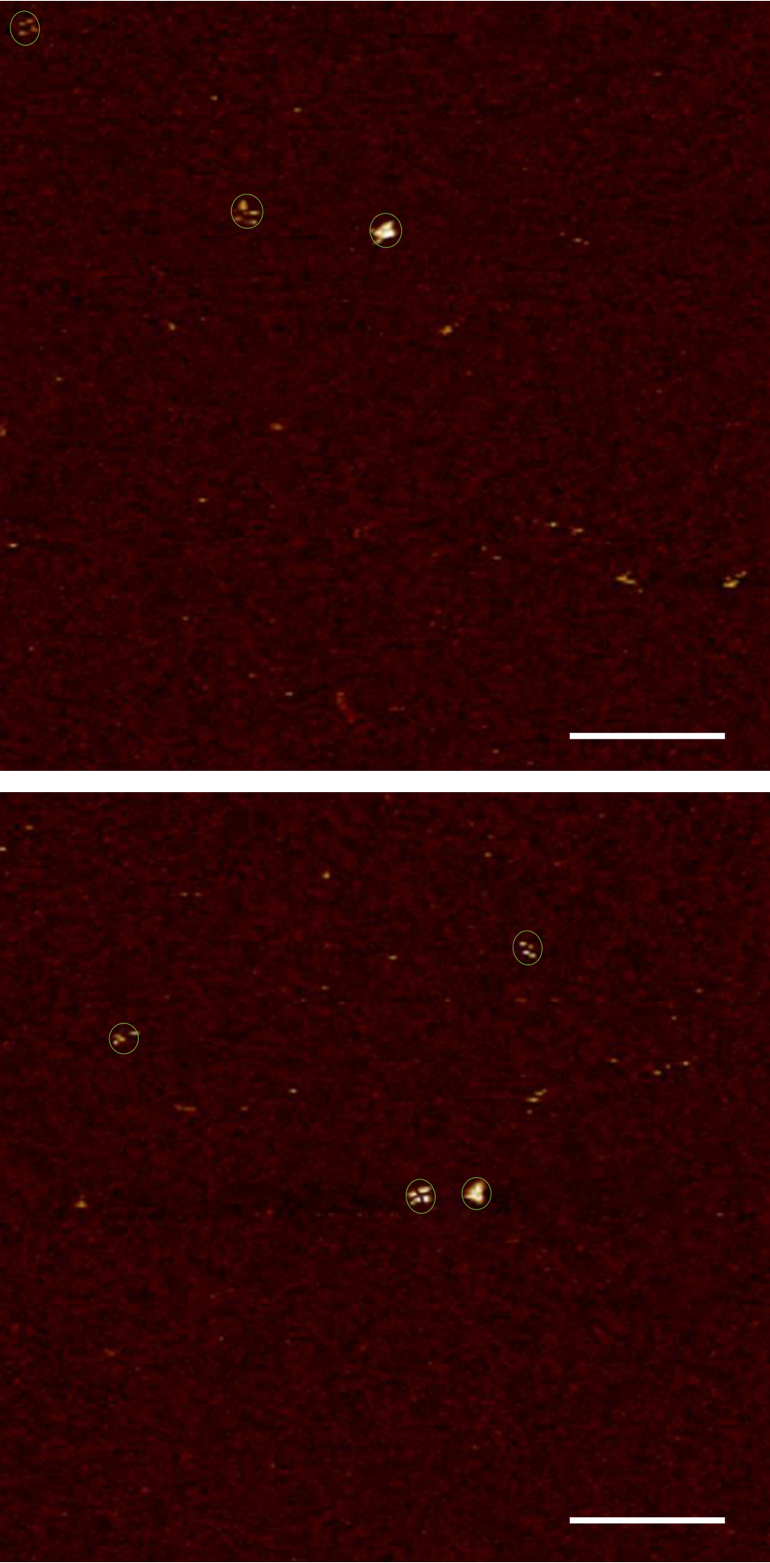

**Figure S11: Wide-field AFM images of recovered square tetramers.** Squares are in green circles. Scale bars: 200nm.

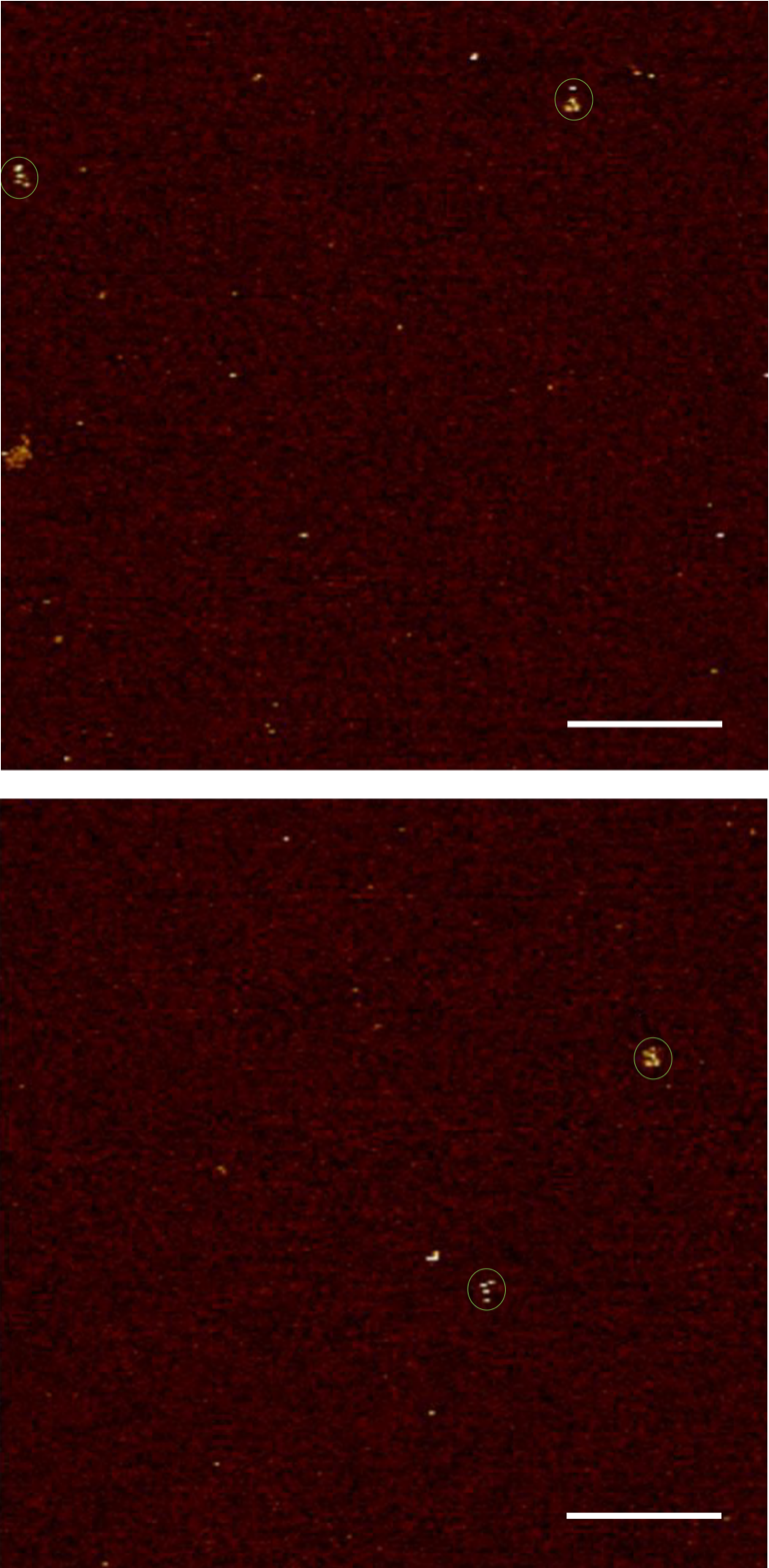

**Figure S12: Wide-field AFM images of recovered Y-shape tetramers.** Y-shaped tetramers are in green circles. Scale bars: 200nm.

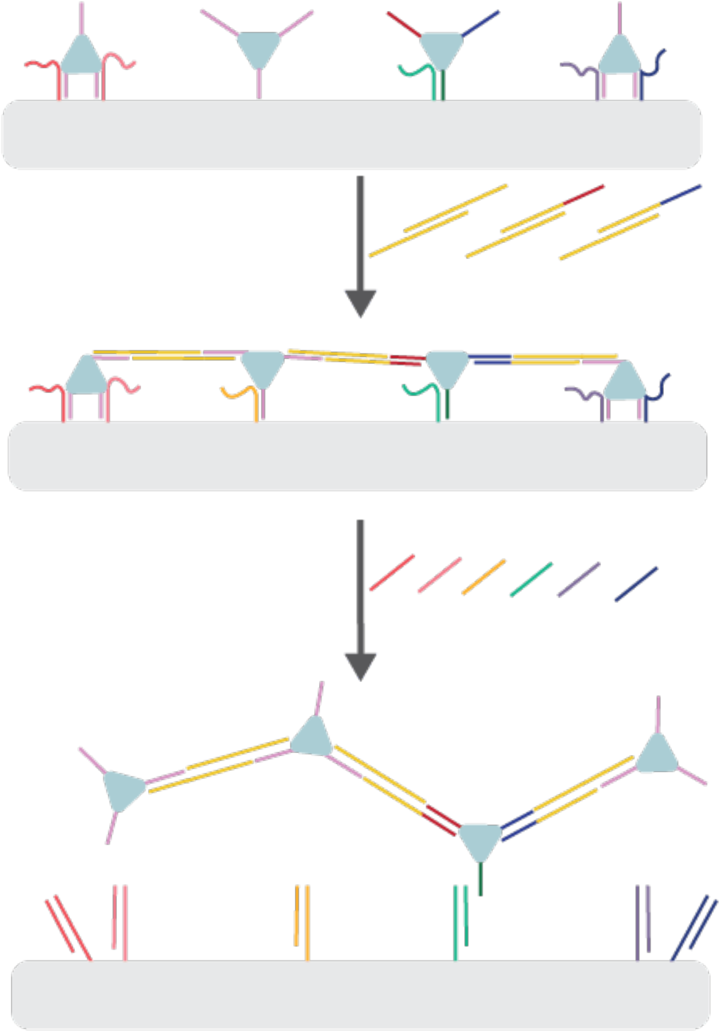

**Figure S13: Assembly can be simplified by using different building blocks.** For example, if an ald_3_ c modified with three different ssDNA strands is included. Three different connectors can be added to connect four ald_3_ building blocks in a single step to produce linear a tetramer.

**Sequence of pscaf1800**

AATAGTGGACTCTTGTTCCAAACTGGAACAACACTCAACCCTATCTCGGGCTATTCTTTTGATTTATAAGGGATTTTGCCGATTTCGGGGTACCTACGAAGAGTTCCAGCAGGGATTCCAAGAAATGGCCAATGAAGATTGGATCACCTTTCGCACTAAGACCTACTTGTTTGAGGAGTGCCTGATGAATTGGCACGACCGCCTCAGGAAAGTGGAGGAGCATTCTGTGATGACTGTCAAGCTCCAATCTGAGGTGGGCAAATATAAGATTGTTATCCCTATCTAGAAGTACGTCCGCGGAGAACACCTGCCACCCGATCACTGGCTGGATCTGTTACGCTTGCTGGGTCTGCCTCGCGGCACATCTCTGGAGAAACTGCTGTTCGGTGACCTGCTGAGAGTTGCCGATACCATCGTGGCCAAGGCTGCTAACCTGAAAGATCTGAACTCACGCGGCCAGGGTGAAGTGACCATCCGCGAATAACTCAGGGAACTGGATTTGTGGGGCGTGGGTGCTGTGTTCACACTGATCGGCTATGAGGACTCCCAGAGCCGCACCTAGAAGCTGATCAAGGATTGGAAGGAGCTCGTCAACCAGGTGGGCGACAATATATGCCTCCTGCAGTCCTTGAAGGACTCACCATACTATAAAGGCTTTGAAGACAAGGTCAGCATCTGGGCAAGGAAACTCGCCGAACTGGACGATAATTTGCAGAACCTCAACCATATTCGCAGAAAGTGGGTTTACCTCGAACCATACTTTGGTCGCGGAGCCCTGCCCAAAGAGCAGACCAGATTCAACAGGGTGGGCGAAGATTTCCGCAGCATCATGACATATATCAAGAAGGACAATCGCGTCACGCCCTTGACTACCCACGCAGGCATTCTAAACTCACTGCTGACCATCCTGGACCAATTGCAGAGATGCCAGCGCAGCCTCAACGAGTTCCTGGAGGCGAAGCGCAGCGCCTTCCCTCGCTTTAACTTCATCGGAGACGATGACCTGCGCGAGATCTTGGGCCAGTCAACCAATTAATCCGTGATTCAGTCTCACCTCAAGAAGCTGTTTGCTGGTATCAACTCTGGCTGTTTCGATGAGAAGTCTAAGCACTATACTGCAATGAAGTCCTTGGAGGGGCAAGTTGTGCCATTCAAGAATAACGTACCCTTGTCCAATAACGTCGAAACCTGGCTGAACGATCTGGCCCTGGAGATGAAGAAGACCCTGGAGGCGCTGCTGAAGGAGTGCGTGACAACTAGACGCAGCTCTCAGGGAGCTGTGGGCCCTTCTCTGTTCCCATCACAGATCTAGTGCTTGGCCGAACAGATCAAGTTTACCGAAGATGTGGAGAACGCAATTAAAGATCACTCCCTGCACCAGATTGAGTAACAGCTGGTGAACAAATTGGAGCAGTATACTAACATCGACACATCTTCCGTAGACCCAGGTAACACAGAGTCCGGTATTCTGGAGCTGAAACTGAAAGCACTGATTCTCGACGGATCCACGCGCCCTGTAGCGGCGCATTAAGCGCGGCGGGTGTGGTGGTTACGCGCAGCGTGACCGCTACACTTGCCAGCGCCCTAGCGCCCGCTCCTTTCGCTTTCTTCCCTTCCTTTCTCGCCACGTTCGCCGGCTTTCCCCGTCAAGCTCTAAATCGGGGGCTCCCTTTAGGGTTCCGATTTAGTGCTTTACGGCACCTCGACCCCAAAAAACTTGATTTGGGTGATGGTTCACGTAGTGGGCCATCGCCCTGATAGACGGTTTTTCGCCCTTTGACGTTGGAGTCCACGTTCTTT

**Table S3: Staple strands for 4HB**

| 3’ End | Sequence |
| --- | --- |
| 0[16] | GTAAAGCAAAGGGAGCGCAAGCGT |
| 0[48] | ACCCAAATAAGCCGGCTTTCTCCA |
| 0[80] | CACCTCAGATTGGAGCCGATGGTATCGGCAAC |
| 0[112] | TGGACTCCCAAGTGTACTTTCAGG |
| 0[144] | TAACCACCACACCCGCGGTCACTT |
| 0[152] | CAATTCATCAGGCACTTCGCGGAT |
| 0[176] | GAATAGCCCGCGTGGACACAAATC |
| 0[200] | CCGAAATCTTTCAGCTAGCCGATCAGTGTGAA |
| 0[240] | CCTTCCAAAGCGAGGGTTAATTGGTTGACTGGCCCAAGATCTGGGTCTCAGCTTCT |
| 0[264] | GAGACTGATACTGCTCACCTGGTTGACGAGCT |
| 0[304] | TAGTATGGTTGGTCCATCTCATCGAAACAGCCAGAGTTGAAATCTGGTTCAAGGAC |
| 0[328] | GTATAGTGTCTCCACAGTCTTCAAAGCCTTTA |
| 0[368] | CAAATTATTTGTCCTTACGTTATTCTTGAATGGCACAACTGCCAAGCATCGGCGAG |
| 0[392] | GTTATTGGAGGGCCCATATGGTTGAGGTTCTG |
| 1[55] | TAGAGCTTGACGGGGACAAGTTTTGGACGTACTTCTAGAT |
| 1[79] | TCTCAGCAGGTCACCGCAATCTTATATTTGCC |
| 1[119] | GGCGCTAGGGCGCTGGAACGTCAACATCACAGAATGCTCC |
| 1[143] | AGTTTGGAACAAGAGTGGTCGTGC |
| 1[175] | CGCGCTTAATGCGCCGCTACAGGGCGAGATAGAAGTAGGT |
| 1[247] | CCGGACTCTGTGTTACCTCGCGCAGGTCATCGTCTCCGAT |
| 1[311] | TCACCAGCTGTTACTCTACCAGCAGAGGCTGCGCTGGCAT |
| 1[375] | AAACTTGATCTGTTCGTGCCCCTCTGGGTAGTCAAGGGCG |
| 1[439] | TGAGAGCTGCGTCTAGCAGATCGTCGCCCACCCTGTTGAA |
| 2[24] | GAGATGTGCCGCGAGGTTCTCCGCTTGGGGTCGAGGTGCC |
| 2[56] | CACTACGTGAACCATC |
| 2[72] | AAACCGTCTATCAGGGCGATGGCC |
| 2[96] | TTAGCAGCCTTGGCCATTGACAGTAGGGCGAA |
| 2[120] | CACCCTGGCCGCGTGACCTGAGGCCCACTATTAAAGAACG |
| 2[152] | CAGTTCCCTGAGTTATCCTCAAACGGTTGAGTGTTGTTCC |
| 2[184] | CACAGCACATCCAATCCCCTTATAAATCAAAA |
| 2[216] | AGGTGCGGCTCTGGGACTGCTGGAACTCTTCGTAGGTACC |
| 2[280] | TGCAGGAGGCATATATAACTCGTTAACAGCTTCTTGAGGT |
| 2[344] | TTTCCTTGCCCAGATGTGCCTGCGCAAGGACTTCATTGCA |
| 2[408] | CGAGGTAAACCCACTTGAAATCTTTCAGCCAGGTTTCGAC |
| 3[31] | GATCGGGTGGCAGGTGCAGACCCACCCCGATT |
| 3[79] | AGGGATAAAACAGCAGGAACGTGGCGAGAAAGGAAGGGAA |
| 3[95] | GAAAGCGAAAGGAGCG |
| 3[135] | TCCACTTTGTTCAGATGCGGTCACGCTGCGCG |
| 3[207] | CTTAGTGCGAAAGGTGCCACGCCCTCCGTCGAGAATCAGTGCTTTCAG |
| 3[223] | GGCAAAATTTCATTGGCCATTTCTTGGAATCCGTCCTCATCCAGAATA |
| 3[271] | GAAGTTAATCCTTGATACGGAAGATGTGTCGATGTTAGTA |
| 3[287] | ATCACGGAAAGGCGCTGCGCTTCGCCTCCAGGTGTCGCCCCAATTTGT |
| 3[335] | CTCTGCAATGAGTCCTGCAGGGAGTGATCTTTAATTGCGT |
| 3[351] | CTTAGACTGGATGGTCAGCAGTGAGTTTAGAACTGACCTTTCTTCGGT |
| 3[399] | TGACGCGACGTCCAGTCTAGATCTGTGATGGGAACAGAGA |
| 3[415] | ACAAGGGTCTTGATATATGTCATGATGCTGCGTCTGCGAACAGCTCCC |

**Table S4. Sequence of sticky-end strands for forming 4HB dimer**

| sticky 0 | GACTATAGTCCAGGGCTCCGCGACCAAAGTATGG |
| --- | --- |
| sticky 1 | TCTGGTCTGCTCTTTGAAGTCAGGCA |
| sticky 2 | ATGGCGCCATTCCAGGGTCTTCTTCATCTCCAGG |
| sticky 3 | TTGTCACGCACTCCTTCAGCAGCGTGCCTGACTT |

**Table S5. Sequences of Attachment strands and Releasing strands (Ra1-Rd2)**

| a1 | CACCTCAGATTGGAGCCGATGGTATCGGCAACTTACCTGACGGAACTCACCGCGCCCCAGCGGGCTAGG |
| --- | --- |
| a2 | TCTCAGCAGGTCACCGCAATCTTATATTTGCCTTACCTGACGGAACTCAGCACGGCGCCGGAGCCTGCC |
| b1 | TGGACTCCCAAGTGTACTTTCAGGTTACCTGACGGAACTCACCCAGGCGCCCGGGGCGGCG |
| c1 | TAACCACCACACCCGCGGTCACTTTTACCTGACGGAACTCATCCGCTACGACTTCCGGGTC |
| c2 | AGTTTGGAACAAGAGTGGTCGTGCTTACCTGACGGAACTCACGGCGTGTCCGCGCTCGCGC |
| d1 | GAATAGCCCGCGTGGACACAAATCTTACCTGACGGAACTCACATACGCGCGCAAGGCCGGG |
| d2 | CGCGCTTAATGCGCCGCTACAGGGCGAGATAGAAGTAGGTTTACCTGACGGAACTCACCGGTGCGTAGCATGGTCAC |
| d1 fill | GAATAGCCCGCGTGGACACAAATC |
| d2 fill | CGCGCTTAATGCGCCGCTACAGGGCGAGATAGAAGTAGGT |
| Ra1 | CCTAGCCCGCTGGGGCGCGGTGAGTTCCGT |
| Ra2 | GGCAGGCTCCGGCGCCGTGCTGAGTTCCGT |
| Rb1 | CGCCGCCCCGGGCGCCTGGGTGAGTTCCGT |
| Rc1 | GACCCGGAAGTCGTAGCGGATGAGTTCCGT |
| Rc2 | GCGCGAGCGCGGACACGCCGTGAGTTCCGT |
| Rd1 | CCCGGCCTTGCGCGCGTATGTGAGTTCCGT |
| Rd2 | GTGACCATGCTACGCACCGGTGAGTTCCGT |

**Table S6. Sequences of Connector strands**

| connector 1a | TTAAGAACCTCTCCG**GAGCAGACCTGACGGAACTCA** |
| --- | --- |
| connector 1b | CGGAGAGGTTCTTAA**GAGCAGACCTGACGGAACTCA** |
| connector 2a | TTAAGAACCTCTCCGTCGTCTGGTATAG**GAGCAGACCTGACGGAACTCA** |
| connector 2b | CTATACCAGACGACGGAGAGGTTCTTAA**GAGCAGACCTGACGGAACTCA** |
| connector 3a | AAGAACCTCTCCGTCGTCTGGTATAGATGTGAATGATG**GAGCAGACCTGACGGAACTCA** |
| connector 3b | CATCATTCACATCTATACCAGACGACGGAGAGGTTCTT**GAGCAGACCTGACGGAACTCA** |
| connector 4a | TTAAGAACCTCTCCG**ACCTGACGGAACTCA** |
| connector 4b | CGGAGAGGTTCTTAA**ACCTGACGGAACTCA** |
| connector 5a | TCGGCGCGATAGGCCGTTAGAG**ACCTGACGGAACTCA** |
| connector 5b | CTCTAACGGCCTATCGCGCCGA**ACCTGACGGAACTCA** |

Note: The original sequence of ssDNA on the ald-ssDNA was 5' TGAGTTCCGTCAGGTCTGCTCT after decreasing the length from 21nt to 15nt, the sequence was changed to 5' TGAGTTCCGTCAGGT.
